# R-loop imbalance compromises virulence in *Salmonella enterica*

**DOI:** 10.1101/2025.01.19.633779

**Authors:** Diego A. Pedraza-Delgado, Julia Jiménez-Espadafor, Miriam Ortiz-Padilla, Joaquín Bernal-Bayard, Francisco Ramos-Morales, Roberto Balbontín

## Abstract

Bacterial virulence and antibiotic resistance are interconnected global threats that can synergize to multiply their adverse effects on health and economy. Two proposed strategies to address this dual challenge are non-biocidal inhibition of virulence traits (anti-virulence strategy) and manipulation of eco-evolutionary dynamics in bacterial populations to hinder dissemination of antibiotic resistance (anti-resistance strategy). Both strategies require identifying factors involved in bacterial gene expression, fitness and evolution. R-loops strongly influence these traits, as well as fitness of bacteria carrying antibiotic resistance mutations. This makes the enzyme responsible for R-loop degradation, the RNase HI, a promising target for anti-resistance approaches. Interestingly, the involvement of R-loops in gene expression could also make RNase HI a potential target for anti-virulence strategies. In this study, we explored this possibility by investigating the effects of RNase HI deficiency on the pathogenicity of *Salmonella enterica*. We found that the absence of RNase HI alters the expression of genes associated with virulence both at the population and single-cell levels, and both *ex vivo* and during infection of mammalian cells. Furthermore, we observed that RNase HI depletion causes defects in phenotypes associated with virulence, such as motility and biofilm formation. Lack of RNase HI also reduces the ability of *Salmonella* to survive within macrophages. However, lack of RNase HI does not significantly affect the invasion of mammalian epithelial cells or their immune response. Overall, our results demonstrate a pleiotropic influence of R-loops on bacterial virulence, suggesting that RNase HI could be a target for anti-virulence strategies. These findings provide a theoretical framework for the development of dual anti-resistance and anti-virulence interventions based on RNase HI targeting.

## INTRODUCTION

The world is currently entering a so-called post-antibiotic era^1^. With a reduced palette of effective antibiotics, bacterial pathogens may cause an enormous health and economic burden worldwide^2^. For instance, mortality associated to antibiotic-resistant infections is estimated to exceed that of cancer by 2050^3^, and was almost 5 millions in 2019^4^. Moreover, there is growing evidence base linking antibiotic resistance and climate change^5^, increasing the scopes of these global challenges. The capacity of bacteria to cause disease is a critical aspect of the antibiotic resistance crisis. Indeed, antibiotic resistance and bacterial virulence can be conceived as two sides of the same coin, as bacterial pathogens represent a smaller threat when can be treated with effective antibiotics, and antibiotic resistant bacteria pose a lesser challenge if are not able to develop pathogenesis. In addition, antibiotic resistance and bacterial virulence can influence one another, multiplying their combined detrimental effects on global health and economy^6^. Thus, there is an imperative need to develop novel strategies to tackle bacterial pathogens and antibiotic resistance, as well as to address their reciprocal influence^1^.

*Salmonella enterica* is a Gram-negative bacterial species that encompasses over 2500 serovars^7^, able to infect a broad range of hosts, causing from gastroenteritis to systemic infection, depending on the serovar and the host^8^. Many *S. enterica* pathogenic functions are encoded by virulence genes, some clustered in horizontally acquired genomic regions known as *Salmonella* pathogenicity islands (SPIs)^9–12^, which can encode secretion systems able to inject into the host cells bacterial proteins known as effectors^13^. These effectors promote invasion of host cells by bacteria and interfere with host functions in order to facilitate pathogenesis^14^. Two of the most relevant and well-studied SPIs are SPI-1 and SPI-2, which encode two distinct Type III Secretion Systems (T3SS) involved in invasion of the intestinal epithelium and proliferation within mammalian cells^15^, respectively. *Salmonellae* are responsible for a large mortality and morbidity in human populations, as well as for a vast economic loss due to infections of farming animals^16,17^. Thus, novel therapeutical interventions against infections caused by *S. enterica*, particularly by antibiotic-resistant strains, are urgently needed.

Two promising approaches recently proposed to tackle the double threat of bacterial virulence and antibiotic resistance are anti-virulence therapy (non-biocidal suppression of pathogenic traits)^18^ and anti-resistance strategy (selection against resistant bacteria by exploitation of eco-evolutionary dynamics in bacterial populations)^19^. A key aspect for the development of either strategy is identifying specific targets involved in bacterial pathogenesis and/or evolution. Interestingly, bacterial cells contain molecular structures, called R-loops, which influence both regulation of gene expression and bacterial evolution^20–23^. Thus, mechanisms involved in R-loop biology could potentially serve as targets for both anti-virulence and anti-resistance therapies.

R-loops are RNA-DNA hybrids formed during transcription, upon the invasion of the DNA template strand by the nascent RNA^24^, which participate in key physiological processes, such as transcription initiation and termination, regulation of gene expression, DNA repair, etc^20–23^. However, if elevated, R-loops can cause transcription stalling, blocks to replication fork progression, inhibition of DNA repair, replication-transcription conflicts and double-strand DNA breaks^25^. Thus, R-loops need to be maintained within an optimal range to remain harmless for the cell^26–28^. RNase HI, whose function is conserved along the phylogenetic tree, is one of the key factors involved in R-loop homeostasis, as it specifically degrades RNA in R-loops^29,30^.

We recently showed that depleting RNase HI function (either genetically or chemically) strongly reduces the fitness of *Escherichia coli* carrying mutations conferring resistance to rifampicin and/or streptomycin^31^. Indeed, lack of RNase HI leads to extinction of resistant clones when competing against susceptible bacteria even at high initial frequency of resistance, abolishing compensatory evolution, both *ex vivo* and in the mammalian gut^31^. Interestingly, depletion of RNase HI can enhance an antimycobacterial treatment^32^, confirming RNase HI as a promising target for anti-resistance approaches.

The involvement of R-loops in transcription^33,34^, and therefore in gene expression, makes possible that R-loop imbalance caused by lack of RNase HI affects expression of virulence genes, potentially altering the capacity of bacterial pathogens to timely develop the program of gene expression that leads to efficient pathogenesis. Indeed, other ribonucleases have been shown to play central roles in the control of bacterial virulence^35,36^, including regulation of SPIs in *Salmonella enterica*^37,38^.

Thus, RNase HI function, which is susceptible to chemical inhibition^31^, could potentially constitute a convergent target for both anti-resistance and anti-virulence approaches that remained unexplored to date. Here we investigated the involvement of R-loops in *S. enterica* virulence, and found that expression of genes associated with pathogenesis are significantly altered in the absence of RNase HI, both at the population and single-cell levels. We also demonstrate that phenotypes related to virulence, such as biofilm formation, motility and growth in low pH, are reduced in a strain lacking RNase HI. Finally, we showed that, albeit lack of RNase HI does not significantly change *Salmonella*’s ability to invade mammalian epithelial cells not their immune response to the bacterium, it negatively affects its capacity to survive within mammalian macrophages. Taken together, our results indicate that RNase HI represents a promising target for anti-virulence strategies and may constitute a novel dual target for both anti-virulence and anti-resistance approaches.

## RESULTS

### *Expression of genes associated with virulence is altered in* Salmonella *lacking RNase HI*

To study the effects of R-loop imbalance on pathogenesis, we first constructed a *Salmonella* strain carrying a scarless in frame deletion in the gene encoding RNase HI: *rnhA*. The deletion was designed in a way that does not affect a promoter of the downstream gene *dnaQ*, which is located within the coding sequence of *rnhA*^39^, and rendered a shorter version of the RNase HI lacking amino acids 44-153, which encompass three out of the four highly conserved carboxylates that form the so-called DEDD motif comprising the active-site core required for its catalytic activity^30^. We then measured the fitness of the *ΔrnhA* mutant strain and wild-type bacteria by running both growth curves and competitions in Lysogeny Broth (LB)^40^, and observed a small but significant fitness defect in the *ΔrnhA* mutant with respect to wild-type bacteria (Figure S1).

In order to test if R-loop imbalance affects expression of genes important for virulence, we took advantage of a collection of luciferase reporter fusions available in the laboratory^41^, and selected a set of 15 genes involved in processes relevant to virulence, such as SPIs, iron uptake, quorum sensing or Type VI Secretion Systems (Table 1). We measured the expression levels of these genes in wild-type and *ΔrnhA* strains under three different conditions: i) standard laboratory conditions (LB with shaking), ii) conditions of induction of SPI-1 (static cultures in LB with 0.3M of NaCl)^42^, and iii) in a medium that mimics the intracellular milieu of mammalian cells (Low Phosphate and Magnesium medium, LPM) and induces SPI-2^43^. As Figure 1 shows, expression of most genes is significantly altered in the *ΔrnhA* mutant in at least one of the conditions tested (Figure 1A), causing in most cases a negative effect (Figure 1B-D). However, upregulation can be observed in certain genes and conditions; for instance, genes encoding the SPI-2 T3SS effector SseK3, the structural component of the system SsaV, and the enterobactin non-ribosomal peptide synthetase EntF in conditions of SPI-2 induction, and the synthase of the quorum molecule autoinducer-2 LuxS in LB and conditions of SPI-1 induction ( Figure 1A).

**Figure 1.**
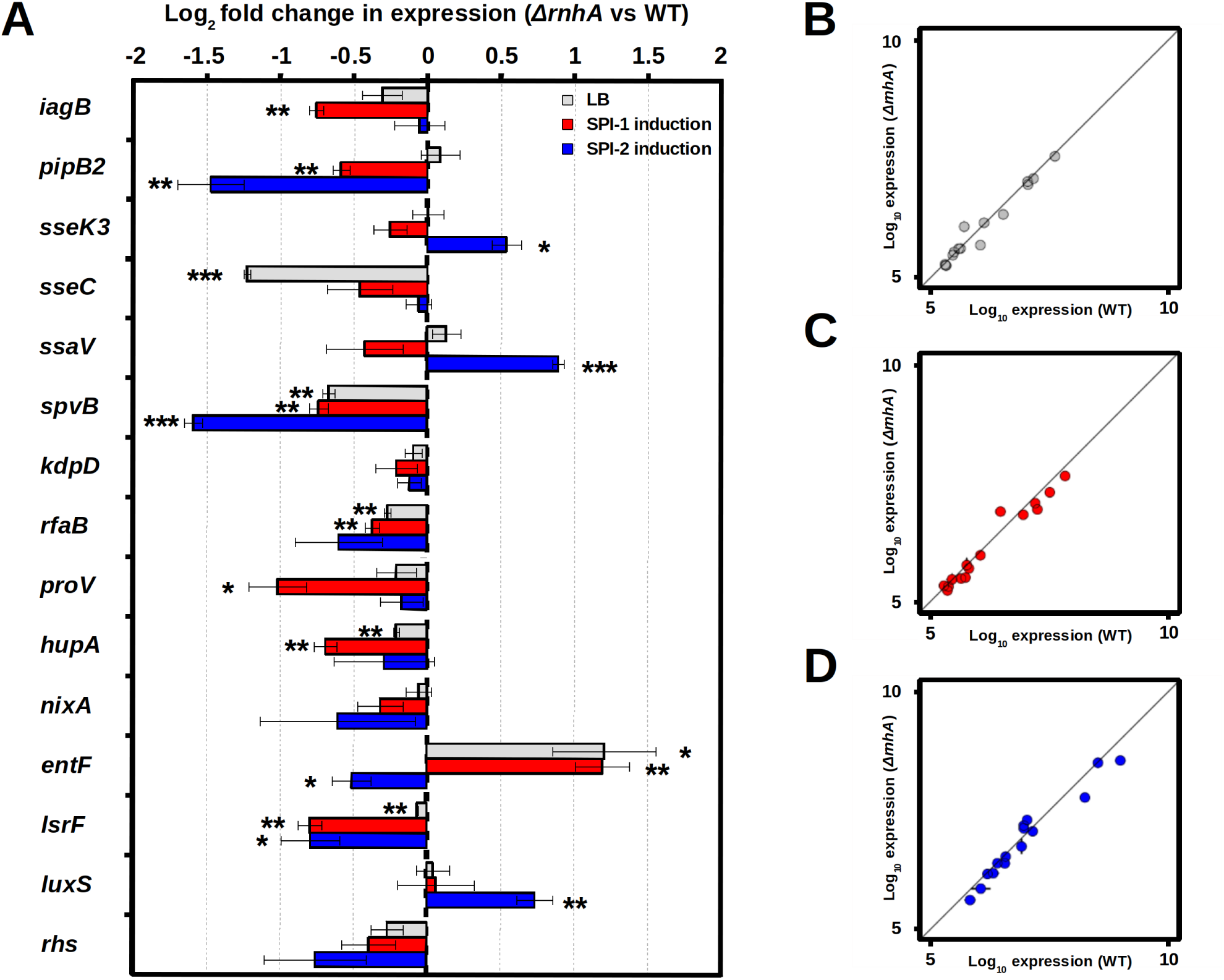
Expression of luciferase reporter fusions to genes associated with *S. enterica* virulence. **A.** Ratio *ΔrnhA*/WT of the area under the curve of luminiscence normalized to OD_600_ of different reporter fusions in LB (grey bars), LB 0.3M NaCl (red bars) or LPM (blue bars). Bars represent average and error bars standard deviation (n = 3 independent biological replicates). N.S. non-significant; *P < 0.05; **P < 0.01; ***P < 0.001; ****P < 0.0001 (one-sample two-tailed Student’s t test). **B-D.** Logarithmic representation of the expression of luciferase reporter fusions in the WT (X axis) and the *ΔrnhA* (Y axis) backgrounds, either in LB (**B**), LB 0.3M NaCl (**C**) or LPM (**D**). The black diagonal line represents the linear regression if expression levels were identical.

**Table 1.** List of genes in the set of luciferase reporter fusions.

| Gene | Location | Function |
| --- | --- | --- |
| *iagB* | SPI-1 | SPI-1 type III secretion system invasion protein |
| *pipB2* | Orphan | SPI-1 and SPI-2 effector |
| *sseK3* | ST64B | SPI-2 effector |
| *sseC* | SPI-2 | SPI-2 effector |
| *ssaV* | SPI-2 | Structural component of the SPI-2 type III secretion system |
| *spvB* | pSLT | Effector |
| *kdpD* | Orphan | Two-component system, sensor histidine kinase |
| *rfaB* | Orphan | Galactosyltransferase |
| *proV* | Orphan | ATP-binding protein of glycine betaine/L-proline ABC transporter |
| *nixA* | Orphan | Putative nickel/cobalt transporter |
| *entF* | Orphan | Enterobactin synthetase |
| *hupA* | Orphan | HU nucleoid-associated protein |
| *lsrF* | Orphan | Quorum Sensing, autoinducer 2 aldolase |
| *luxS* | Orphan | Quorum sensing, autoinducer 2 synthase |
| *rhs* | SPI-6 | Type VI secretion system |

### The absence of RNase HI affects expression of SPI-1 at the single-cell level

Expression of the SPI-1 is not homogeneous across the entire population of *S. enterica*. Instead, it shows bistability; that is, genes in the SPI-1 are expressed only in a subset of the total population, whereas another fraction of the population does not express them^44^. Since lack of RNase HI affects expression of a SPI-1 gene (*iagB*), we next tested if RNAse HI depletion influences expression of SPI-1 at the single-cell level. We constructed wild-type and *ΔrnhA* strains carrying: i) a copy of the gene encoding the superfolder green fluorescent protein (*sfGFP* hereafter) under the control of the promoter and ribosome binding site (RBS) of the SPI-1 gene *sicA,* inserted in the pseudogene locus *malYX*, and ii) a copy of the gene encoding the *mCherry* red fluorescent protein under the control of a constitutive promoter, inserted in the pseudogene STM14_2013. These strains allowed us to distinguish viable bacteria in a regular transcriptional status (those expressing *mCherry*) from compromised or dead cells (those not expressing it), and to simultaneously measure by flow cytometry percentages of viable bacteria expressing or not the SPI-1 (bacteria either showing both red and green fluorescences or only red fluorescence, respectively) in each genetic background (Figure 2A). Interestingly, we observed that lack of RNase HI significantly affects the percentage of bacteria that expresses SPI-1 both in rich (Figure 2B, top panel) and minimal medium (Figure 2B, bottom panel), demonstrating that R-loop imbalance indeed influences bistability in the expression of SPI-1.

**Figure 2.**
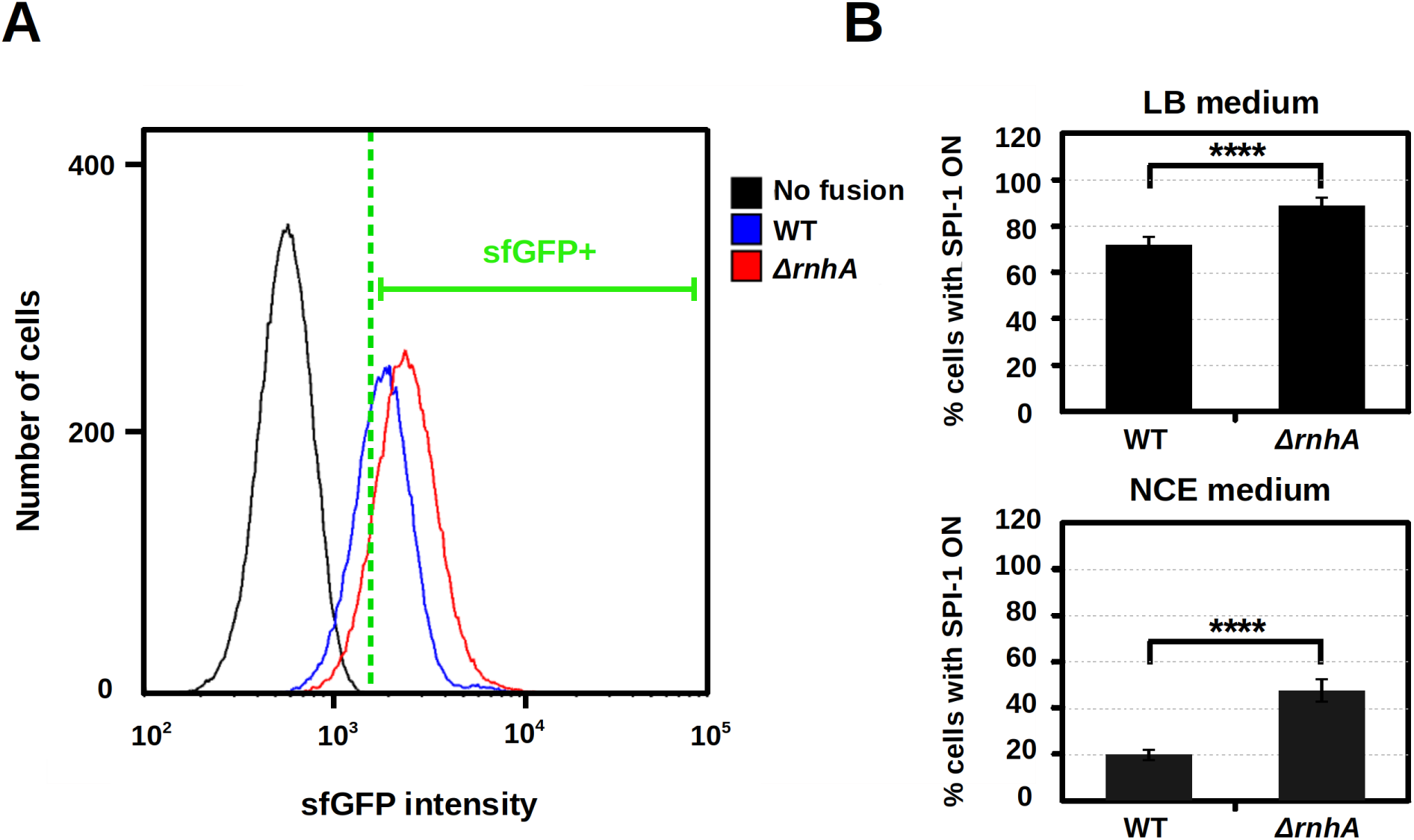
Lack of RNase HI affects fluorescence intensity of bacteria carrying the *P_sicA_-sfGFP* fusion. **A.** Distribution of sfGFP fluorescence intensity in populations of wild-type (blue line) and *ΔrnhA* (red line) bacteria carrying the *P_sicA_-sfGFP* fusion, and in a control not carrying the fusion (black line), in LB. The vertical green dashed line represents the threshold of fluorescent versus non-fluorescent bacteria, determined using non-fluorescent bacteria as control. **B.** Percentage of cells with the *P_sicA_-sfGFP* fusion induced in wild-type and *ΔrnhA* bacteria, either in LB (top panel) or in NCE (bottom panel) media. Bars represent average and error bars standard deviation (n = 6 independent biological replicates). N.S. non-significant; *P < 0.05; **P < 0.01; ***P < 0.001; ****P < 0.0001 (two-tailed Student’s t test).

The results described above confirm that lack of RNase HI alters expression of genes encoding proteins directly involved in virulence, such as T3SS effectors, but also influences genes coding for proteins involved in mechanisms relevant for bacterial physiology that can influence pathogenesis, such as synthesis of enterobactin, signal transduction, and synthesis and processing of the quorum sensing molecule autoinducer-2^45–47^. Particularly, some of these regulatory circuits can influence phenotypes previously connected with bacterial pathogenesis. For instance, quorum sensing has been shown to affect both biofilm formation, bacterial motility and virulence^48,49^. Conversely, SPI-1 has been reported to modulate *Salmonella* motility^50^. This intricate cross-regulation opens the possibility that RNase HI depletion pleiotropically affects multiple phenotypes associated with virulence.

### RNase HI depletion negatively influences biofilm formation and bacterial motility

Biofilms play a relevant role in pathogenicity of *S. enterica*^51^. We thus first used the well-established colony morphology assay^52^, to measure the ability to form biofilms of the *ΔrnhA* mutant, using wild-type bacteria and a mutant lacking the master regulator of biofilm formation CsgD (*ΔcsgD*) as controls. Figure 3A shows that the strain lacking RNase HI is able to form wrinkly structures similar to those of the wild type, and different from the smooth colonies formed by the *ΔcsgD* control. However, colonies of *ΔrnhA* are unable to expand radially as wild-type colonies do, presenting a somewhat intermediate phenotype, not as smooth as the *ΔcsgD* control but not as structured and expanded as wild-type bacteria. In order to quantify these differences, we compared the same genotypes using the crystal violet assay^53,54^. This allowed us to ratify that the mutant lacking RNase HI is not able to form biofilms as efficiently as wild-type bacteria, but still forms better biofilms than the *ΔcsgD* control (Figure 3B). Using controls as references, the ability to form biofilms in the *ΔrnhA* mutant was estimated to be reduced approximately 65% with respect to the wild type (Figure 3C).

**Figure 3.**
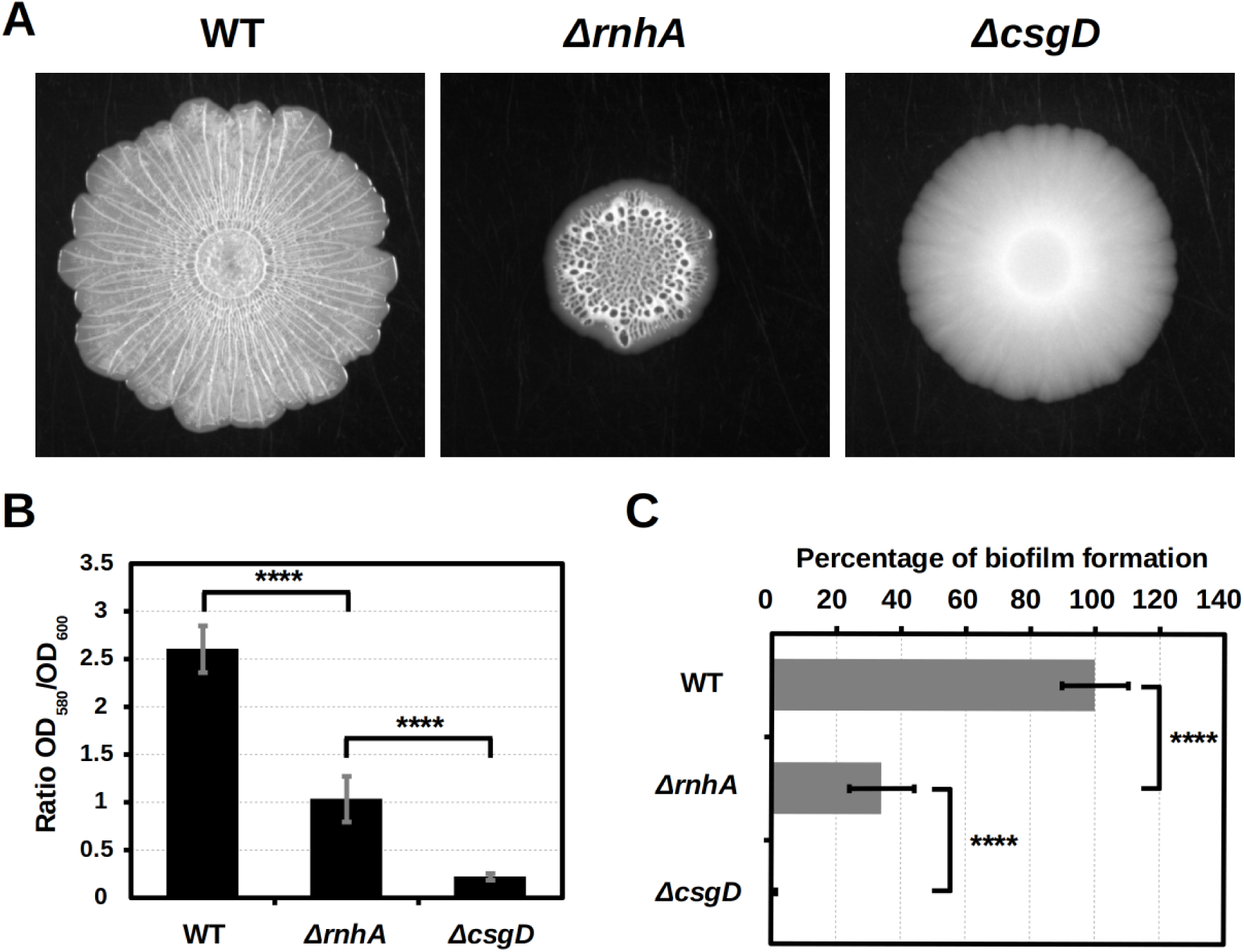
Lack of RNase HI causes reduction in biofilm formation. **A.** Colony morphology of wild-type bacteria (left panel), the *ΔrnhA* mutant (center panel) and the *ΔcsgD* mutant, unable to form biofilms (right panel), after 7 days of incubation at 30 °C in LB agar without salt. **B.** Ratio between the OD_580_ (measuring concentration of crystal violet) and OD_600_ (measuring cell density) of wild-type bacteria, the *ΔrnhA* mutant and the *ΔcsgD* mutant, unable to form biofilms, after 48 h of incubation at 25 °C in LB without salt. **C.** Percentage of biofilm formation of the *ΔrnhA* mutant with respect to wild-type bacteria (considered to form 100%) and the *ΔcsgD* mutant (considered to form 0%). Bars represent average and error bars standard deviation (n = 9 independent biological replicates). N.S. non-significant; *P < 0.05; **P < 0.01; ***P < 0.001; ****P < 0.0001 (two-tailed Student’s t test).

As bacterial motility is also linked to *Salmonella*’s virulence^55^, and connected with some of the genes whose expression is altered by RNase HI depletion, we next tested the effects of lack of RNase HI on it, using the classic bacterial motility assay^56^. We compared motility of bacteria lacking RNase HI with those of the wild type and a non-flagellated *ΔfliGHI* mutant^57^, as controls, and found that the mutant lacking RNase HI is significantly less motile than the wild type, albeit not completely non-motile, as it is the *ΔfliGHI* mutant (Figure 4A). According to controls, the *ΔrnhA* mutant shows a reduction of around 40% in motility compared to wild-type bacteria (Figure 4B).

**Figure 4.**
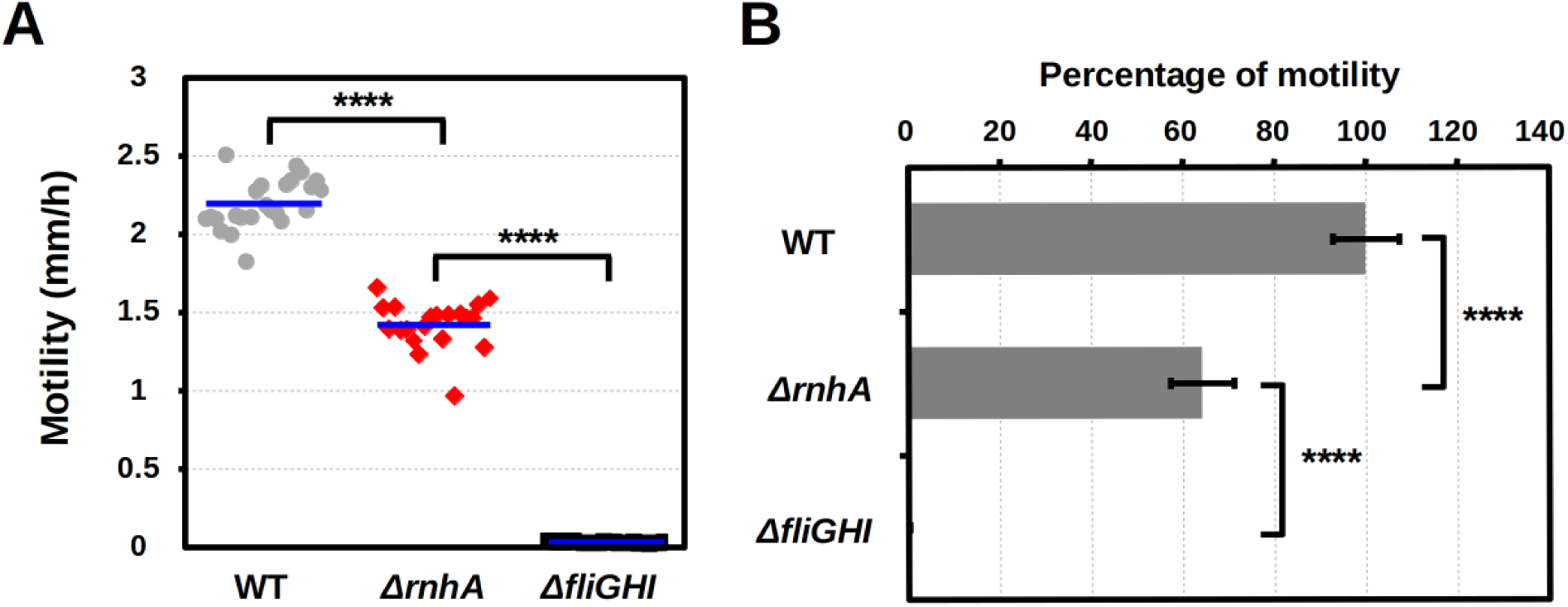
RNase HI depletion negatively affects bacterial motility. **A.** Motility (expressed in milimeters per hour) of wild-type bacteria, the *ΔrnhA* mutant, and the non-motile *ΔfliGHI* mutant. Blue lines represent mean values. **B.** Percentage of motility of the *ΔrnhA* mutant with respect to wild-type bacteria (considered to move 100%) and the *ΔfliGHI* mutant (considered to move 0%). Bars represent average and error bars standard deviation (n ≥ 20 independent biological replicates). N.S. non-significant; *P < 0.05; **P < 0.01; ***P < 0.001; ****P < 0.0001 (two-tailed Student’s t test).

### Expression of SPI-2 is altered in the ΔrnhA mutant during infection of mammalian epithelial cells

In order to test if the influence of R-loops on expression of virulence genes also occurs *in vivo*, we then constructed wild-type and *ΔrnhA* strains carrying the *P_sicA_-sfGFP* reporter fusion of the SPI-1 described above and also an additional reporter fusion, *P_ssaB_-mCherry*, in which the *mCherry* is under the control of the promoter and RBS of the SPI-2 gene *ssaB,* inserted in the pseudogene locus *STM14_2013.* These double reporter strains allow to simultaneously measure expression of both SPI-1 and SPI-2 by flow cytometry using wild-type and *ΔrnhA* strains carrying constitutively expressed *sfGFP* or *mCherry* as controls. We infected HeLa human epithelial cells with these wild-type and *ΔrnhA* strains carrying the double reporter for SPI-1 and SPI-2 or constitutively expressing *sfGFP* and *mCherry,* and analyzed the number of bacterial cells expressing each island along the infection. Strains constitutively expressing each fluorescence allowed us: i) to monitor progression of the infection in each genetic background and ii) to set up a threshold to distinguish fluorescent from non-fluorescent bacterial cells, and therefore which ones can be considered to show SPI-1 and/or SPI-2 activation. Interestingly, initially both genotypes seem equally able to invade host cells, but differences in number of bacteria can be observed as the infection progresses (Figure 5A), suggesting that mid-term survival of the strain lacking RNase HI may be compromised. Regarding expression of SPIs, we observed similar expression patterns in the wild type and the mutant lacking RNase HI at 30 minutes after infection. However, at 24 hours the *ΔrnhA* shows a significantly greater percentage of cells expressing SPI-2 than wild-type bacteria (Figure 5B).

**Figure 5.**
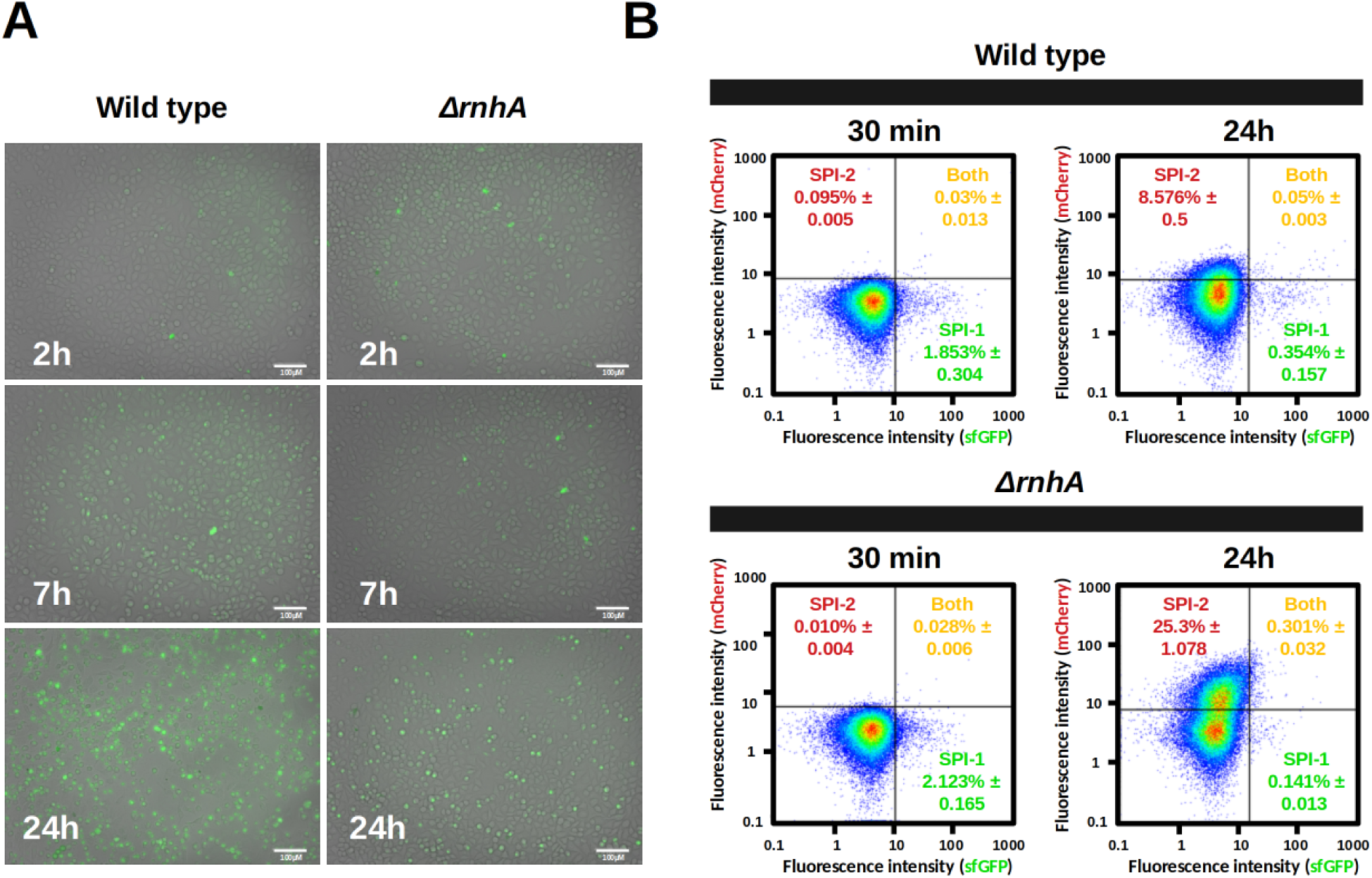
Lack of RNase HI affects expression of a *P_ssaB_-mCherry* fusion during infection of epithelial cells. **A.** Fluorescence microscopy image of HeLa cells infected with wild-type (left panels) or *ΔrnhA* (right panels) bacteria constitutively expressing sfGFP at 2, 7 and 24 h after infection. **B.** Distribution of sfGFP and mCherry fluorescence intensity in wild-type (top panel) and *ΔrnhA* (bottom panel) bacteria carrying the *P_sicA_-sfGFP* and *P_ssaB_-mCherry* fusions, within HeLa epithelial cells at 30 minutes and 24 h after infection. Thresholds for sfGFP and mCherry fluorescences were determined using strains expressing either fluorescence constitutively and non-fluorescent control strains. Percentages represent the average of two independent experiments.

### *Invasion of mammalian epithelial cells by* Salmonella *is not affected by loss of RNase HI*

The fact that lack of RNase HI alters expression of SPI-1, SPI-2, and that of other genes associated with virulence, together with the observed effects on phenotypes relevant for pathogenesis, such as motiliy and biofilm formation, may cause the *ΔrnhA* mutant to be defective in virulence. To test this, we first measured the ability to invade human HeLa human epithelial cells of wild-type bacteria, the *ΔrnhA* mutant, and a double mutant where SPI-1 and SPI-2 are deleted (*ΔSPI-1 ΔSPI-2* hereafter)^58^. Consistent with the results shown in Figure 5, we did not observe any significant difference in invasion by wild-type bacteria or by mutants lacking RNase HI (Figure 6), indicating that R-loop imbalance does not affect the ability of *Salmonella* to invade epithelial cells.

**Figure 6.**
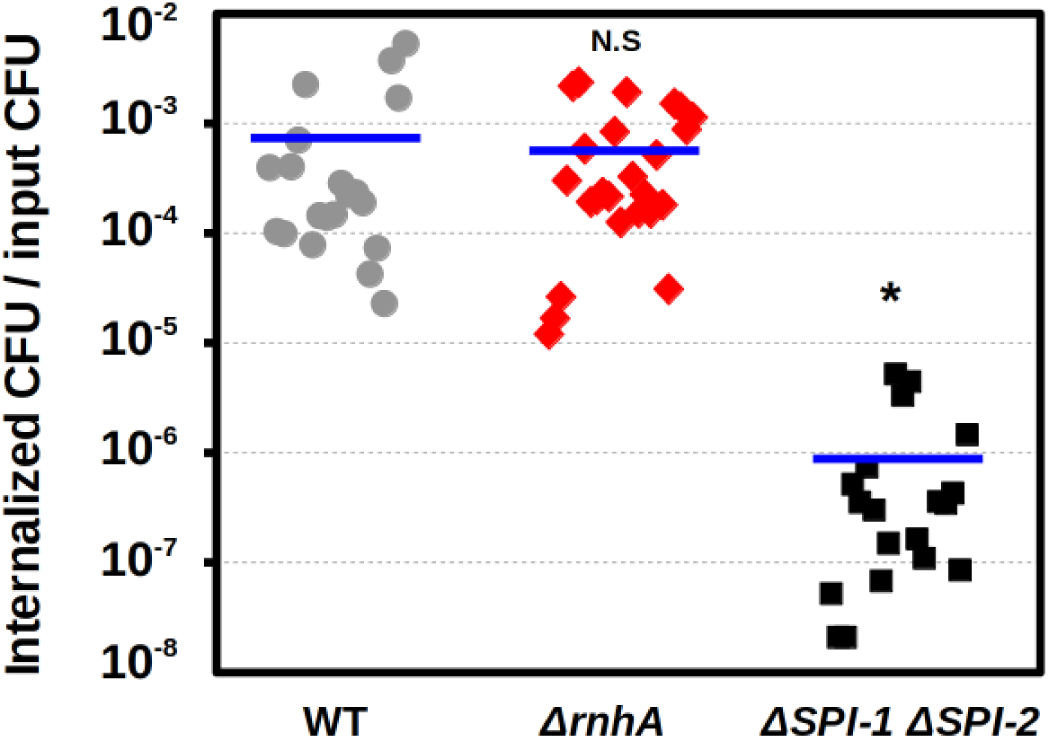
R-loop imbalance does not affect invasion of human epithelial cells. Rate of invasion during individual infection of HeLa human epithelial cells of either wild-type bacteria (grey circles), a mutant lacking the two major *Salmonella* Pathogenicity Islands SPI-1 and SPI-2 (black squares), as controls, and the *ΔrnhA* mutant (red diamonds). Horizontal bars represent the mean (n ≥ 20). N.S. non-significant; * = P < 0.05; ** = P < 0.01; *** = P < 0.001;**** = P < 0.0001 (one-sample two-tailed Student’s t-test).

### Lack of RNase HI negatively affects Salmonella’s ability to proliferate within mammalian macrophages

In order to confirm or refute the seeming decline of mutants without RNase HI over the course of infection (Figure 5), we decided to specifically test potential effects of R-loop imbalance on intracellular proliferation. We thus next performed competitions during infection of RAW264.7 murine macrophages^59^ of either the wild type, the *ΔrnhA* mutant or the *ΔSPI-1 ΔSPI-2* mutant against a reference wild-type strain labeled with a constitutively expressed *lacZ*. As Figure 7 shows, the capacity of the mutant lacking RNase HI is severely impaired with respect to the wild type, showing proliferation similar to that of the *ΔSPI-1 ΔSPI-2* control.

**Figure 7.**
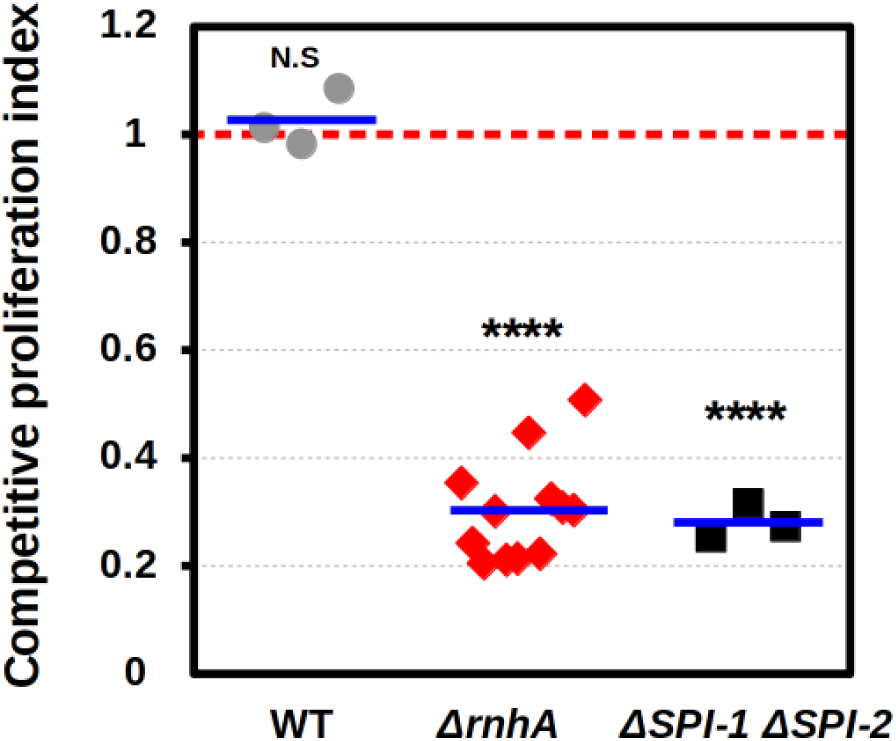
Mutants lacking RNase HI are defective in proliferation within macrophages. Competitive index in proliferation within murine macrophages of either wild-type bacteria (grey circles), a mutant lacking the two major *Salmonella* Pathogenicity Islands SPI-1 and SPI-2 (black squares), as negative control, and the *ΔrnhA* mutant (red diamonds). Horizontal bars represent the mean (n ≥ 3 independent biological replicates). The red dashed line indicates the value of mean competitive proliferation index when both genotypes show similar ability to proliferate. N.S. non-significant; * = P < 0.05; ** = P < 0.01; *** = P < 0.001;**** = P < 0.0001 (one-sample two-tailed Student’s t-test).

### *Secretion of interleukin 8 by mammalian epithelial cells in response to* Salmonella *is not significantly affected by RNase HI depletion*

Infection of host cells by bacterial pathogens can trigger immune responses^60^. In turn, bacterial pathogens have evolved a complex weaponry of virulence factors to evade these immune defenses, at least temporarily^14,61^. Secretion of the pro-inflammatory cytokine interleukin-8 (IL-8) by human epithelial cells has been shown to rapidly increase in response to *S. enterica*^62–65^, and to be downmodulated by the bacterium^66^. In order to test the effects of R-loop imbalance on both cell host immune response and immunomodulation by *Salmonella*, we measured by ELISA levels of IL-8 in the supernatant of HeLa human epithelial cells infected by the *ΔrnhA* mutant, and compared these amounts with those from supernantants of cells infected by either wild-type bacteria or the *ΔSPI-1 ΔSPI-2* mutant, or from non-infected cells, as controls. We did not detect any significant difference in IL-8 secretion between wild-type and *ΔrnhA* bacteria, whereas the *ΔSPI-1 ΔSPI-2* mutant shows IL-8 levels slightly above those in non-infected cells (Figure 8).

**Figure 8.**
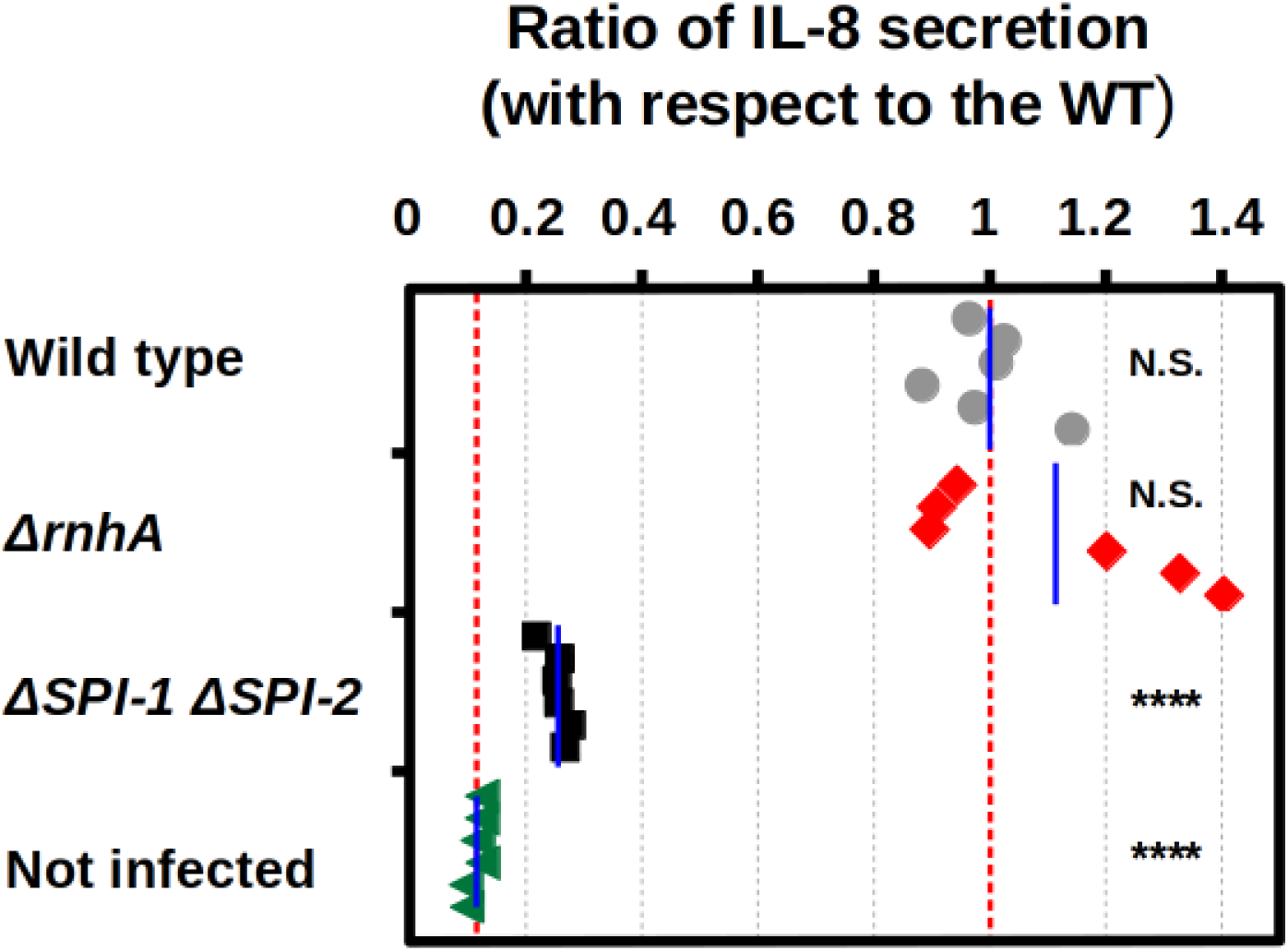
Mutants lacking RNAse HI trigger a normal immune response. Ratio of IL-8 secretion by HeLa epithelial cells at 2 hours after infection by wild-type bacteria (grey circles), a mutant lacking the two major *Salmonella* Pathogenicity Islands SPI-1 and SPI-2 (black squares), as controls, the *ΔrnhA* mutant (red diamonds), and non-infected cells (green triangles). Data are presented as ratios between the corresponding value and the mean values of cells infected by wild-type bacteria. Blue lines represent mean values of six independent biological replicates in each group, with two technical replicates each. Red dashed lines indicate mean levels in non-infected cells and in cells infected by wild-type bacteria. N.S. non-significant; * = P < 0.05; ** = P < 0.01; *** = P < 0.001;**** = P < 0.0001 (one-sample two-tailed Student’s t-test).

## DISCUSSION

The advent of the post-antibiotic era poses the greatest challenge to modern medicine in human history. Numerous medical interventions require efficient antibiotics (e.g. surgery, dental care, treatment of traumatic injuries, etc.), and multiple medical conditions enhance susceptibility to bacterial pathogens (e. g. cancer, AIDS, etc.), making functional antibiotics a necessity to protect the population affected by these conditions^67,68^. Thus, novel strategies to treat bacterial infections and/or prevent the development and dissemination of antibiotic resistances are urgently needed. Anti-resistance and anti-virulence strategies have been proposed as innovative approaches to tackle the increasing threat of antibiotic resistant pathogens^18,19^. We recently reported that the enzyme responsible for degradation of R-loops, the RNase HI, is involved in maintenance of fitness in bacteria carrying antibiotic resistance mutations^31^, being thus a promising candidate as target for anti-resistance approaches. Moreover, the role of R-loops in transcription made plausible that depletion of RNase HI function could affect expression of genes relevant for virulence and therefore cause reduced pathogenicity. We tested this hypothesis by analyzing the effects of lacking RNase HI on phenotypes linked to *Salmonella*’s virulence. We found that, despite not causing a great fitness defect (Figure S1), lack of RNase HI significantly affects expression of multiple genes related to virulence, including genes involved in signal transduction, metal uptake, quorum sensing and genes in SPI-1 and SPI-2 ( Figure 1). These effects on expression of virulence genes also occurs at single-cell level (Figure 2), and during infection of mammalian cells (Figure 5). Moreover, several phenotypes relevant for pathogenicity, such as biofilm formation (Figure 3), and motility (Figure 4), are also negatively affected by RNase HI depletion. This suggests that altered expression of genes associated with virulence caused by the excess of R-loops in the absence of RNase HI renders *Salmonella* unable to properly develop the transcriptional program required for efficiently develop the complete pathogenic course. For instance, reduced motility decreases bacterial ability to swim towards mammalian cells^69^, and defective biofilms affects attachment to mammalian tissues^70^. In addition, excess of R-loops has consequences for bacterial physiology that are particularly disadvantageous during infection. For instance, preliminary results indicate that the mutant lacking RNase HI shows reduced growth at low pH (Figure S2), which is a condition characteristic of the intracellular millieu. Moreover, this strain shows a reduction in proliferation within mammalian macrophages comparable to that of a double SPI-1/SPI-2 deletion mutant (Figure 7). This defect is likely due to misregulation of virulence genes, leading to poor coordination between secretion machinery assembly and effector translocation^71^, as well as to pervasive RNA polymerase backtracking during intracellular growth^72^, which promotes formation of R-loops^33^. Interestingly, invasion of human epithelial cells is not affected in *Salmonella* undergoing R-loop imbalance (Figure 6), nor it is the immune response of epithelial cells to this genotype (Figure 8). Thus, the inability to fully develop pathogenesis while inducing a normal immune response in the host opens the possibility of using RNase HI mutants as live vaccines^73^.

In the onset of a post-antibiotics era, where one of the biggest achievements of modern medicine is under siege, novel approaches to tackle bacterial infections and to reduce antibiotic resistance are desperately needed. In summary, our results demonstrated that R-loop imbalance pleiotropically affects multiple traits important for bacterial pathogenesis, making it a suitable target for anti-virulence strategies. Importantly, as previous results revealed the potential of RNase HI as a target for anti-resistance interventions^31,32^, RNase HI has thus the potential to become a dual target for both anti-virulence and anti-resistance strategies, representing a promising intervention against one of the most pressing challenges to global health.

## MATERIALS AND METHODS

### Bacterial strains, media, and growth conditions

All strains in this study (Table S1) are *Salmonella enterica* subspecies *enterica* serovar Typhimurium strain ATCC 14028s^74^. Mutant strains were constructed by Lambda-Red recombineering^75^ and/or scarless DNA recombineering^76^, followed by transduction to a clean genetic background using P22 HT 105/1 *int201*^77^. Constitutively fluorescent strains harbor a copy of either superfolder green fluorescent protein (*sfGFP*) or mCherry fluorescent protein (*mCherry*), under the control of the P_LTetO-1_ constitutive promoter and a strong RBS^78^, inserted in the pseudogene locus STM14_2013. These constructions were generated by Lambda-Red recombineering using preexisting constructions^79,31^ as templates. Fluorescent reporter fusions were generated by one-step recombineering^80^ and P22 HT 105/1 *int201* transduction. Luciferase fusions were derivatives resulting from P22 HT 105/1 *int201* transduction of fusions obtained in a previous study^41^. Cultures were grown in either Lysogeny Broth^40^ (LB, Lennox formulation), carbon-free minimal medium (NCE)^81^ broth supplemented with 0.4% glucose, LB 0.3M NaCl^42^, or low phosphate, low magnesium-containing medium (LPM)^43^, and incubated at 37°C in either glass inocule tubes with shaking at 200 r.p.m or without shaking, in a INFORs HT Multitron Standard shaker, or in round-bottom 96-well plates with shaking (700 rpm) in a Grant-bio PHMP-4 benchtop incubator. Except for motility and colony morphology assays, solid medium was LB containing 1.5% agar. Media were supplemented when necessary with antibiotics at the following concentrations: ampicillin (100 µg/mL), kanamycin (50 µg/mL), chloramphenicol (10 µg/mL), and tetracycline (20 µg/ mL).

### Growth curves and luciferase expression assays

Strains were streaked individually onto LB agar plates supplemented with the appropriate antibiotic, and incubated overnight at 37 °C. The next day, independent colonies from each strain were inoculated separately in LB supplemented with the appropriate antibiotic (100 µl per well) in a 96-well plate and incubated overnight at 37 °C with shaking (700 rpm). The next day, 1:20 dilutions of the overnight cultures (10µl of culture in 190µl medium) were made, and 10µl of that diluted solution was used to inoculate round-bottom 96-well plates containing 90µl of the corresponding medium. Then plates were incubated at 37 °C with double orbital shaking set as “fast” in a Biotek SYNERGY HTX® multi-mode benchtop microplate reader (Agilent, Santa Clara, CA, United States of America), and OD_600_/luminiscence were measured every 30 min during 24 h.

### Competitive fitness assays

The relative fitness (selection coefficient per generation) analyzed genotype was measured by competitive growth against an isogenic strain carrying a Mu*d*J element (which encodes the *lacZ* gene) inserted in the gene *trg*, which was shown not to affect bacterial virulence^59^, as reference. Competitor strains were first streaked out of their respective frozen vials to LB supplemented with the appropriate antibiotic, then individual colonies were inoculated separately in medium without antibiotics and incubated overnight (approximately 16 h); the next morning, the number of cells in each culture was estimated by OD_600_ and 1:1 mixtures of Lac+ and Lac- bacteria were inoculated in glass tubes and incubated at 37°C with shaking (200 r.p.m) for 24 h. Initial and final frequencies of the strains were estimated by plating dilutions onto plates supplemented with X-gal, and counting Colony Forming Units (CFUs). The number of generations was estimated from that of the reference sensitive strain, and the selection coefficient was determined as described below for each independent competition. The formula used to calculate the selection coefficient was *s*=[ln ( *NSf* / *NRf*) *−* ln ( *NSi* / *NRi*)]/ln ( *NRf* / *NRi*), being NRi and NRf the initial and final number of reference bacteria, and NSi and NSf the initial and final number of the sample bacteria that is being analyzed.

### Flow cytometry

Flow cytometry was performed using either a MACSQuant® Analyzer VYB flow cytometer (Miltenyi Biotec, Bergisch Gladbach, Germany) or a CytoFlex S® flow cytometer (Beckman Coulter, Brea, CA, United States of America), equipped with 488 nm and 561 nm lasers used for scatter parameters and sfGFP and mCherry excitation. Regarding optical configuration for detection, sfGFP was detected using bandpass filters of 525/50 nm (MACSQuant®) or 525/40 nm (CytoFlex S®), and mCherry was detected using bandpass filters of 615/20 nm (MACSQuant®) or 610/20 nm (CytoFlex S®). To detect bacteria, the analyzers were equipped with either a photomultiplier tube (PMT) in the case of the MACSQuant® and an avalanche photodiode (APD) in the case of the CytoFlex S®. Samples were acquired using either MACSQuantify™ 2.13.3 software (MACSQuant®) or CytExpert version 2.3 software (CytoFlex S®), and analyzed using FlowJo version 10.9.0 software (Tree Star, Inc., Ashland, OR, United States of America). Prior to flow cytometry measurement, bacterial cultures were diluted in filtered phosphate-buffered saline (PBS).

### Colony morphology assays

Strains were streaked individually onto LB agar plates supplemented with the appropriate antibiotic, and glass tubes containing LB were inoculated with six independent colonies of each strain and incubated overnight at 37 °C. The next day, 5 µL of each culture were used to inoculate a LB agar without salt (Yeast Extract 5g/L, Tryptone 10g/L)^52^ plate. Images of bacterial colonies after 7 days of incubation at 30°C were acquired using a Gel Doc XR+® Molecular Imager (Bio-Rad, Hercules, CA, United States of America) and Image Lab version 5.1 software.

### Crystal violet assays

Strains were streaked individually onto LB agar plates supplemented with the appropriate antibiotic, and glass tubes containing LB without salt (Yeast Extract 5g/L, Tryptone 10g/L) ^52^ were inoculated with three independent colonies of each strain and incubated overnight at 37 °C. The next day, overnight cultures were diluted 1:100 in LB without salt, six technical replicates of 100 µL from each subculture were placed in wells of a round-bottom 96-well plate to inoculate a LB without salt (Yeast Extract 5g/L, Tryptone 10g/L), and the plate was incubated at 25°C for 48 h, without shaking. Then OD _600_ of the cultures were measured, the medium containing planctonic bacteria was removed, wells were washed three times with sterile PBS, 200 µL of methanol was added, and the plate was incubated 15 minutes at room temperature in static to fix the bacterial biofilm. After the methanol was removed, the plate was left to dry for 10 minutes at room temperature in static, 125 µL of 0.1 crystal violet was added, and the plate was incubated 15 minutes at room temperature in static to stain bacterial biofilms. Then wells were washed again three times with PBS, and the plate was left to dry for at least 2 h. Once dry, 250 µL of 33% acetic acid was added to each well, and the plate was incubated 15 minutes at room temperature in static, to dissolve the crystal violet attached to the biofilms, and then OD_580_ was measured.

### Motility assays

Strains were streaked individually onto LB agar plates supplemented with the appropriate antibiotic, and glass tubes containing LB were inoculated with six independent colonies of each strain and incubated overnight at 37 °C. The next day, a sterile toothpick was soaked in each culture and used to inoculate a motility agar plate (5 g/L NaCl, 10 g/L Tryptone, 3 g/L agar)^56^. Bacterial growth halos were compared after 14 h of incubation at 37°C.

### Infection of HeLa human epithelial cells

HeLa (human epithelial; ECACC no. 93021013) cells were seeded in 24-well plates at 1.5 × 10 ^5^ cells per well and incubated in DMEM supplemented with 10% fetal calf serum for 24 h at 37 °C with 5% CO2. Bacterial strains were streaked individually onto LB agar plates supplemented with the appropriate antibiotic, and glass tubes containing LB were inoculated with independent colonies of each strain and incubated overnight at 37 °C. Then bacteria were added to the cell culture at a multiplicity of infection of 100. To remove noninternalized bacteria, cell cultures were washed twice with sterile PBS 30 minutes after infection, overlaid with DMEM containing 100 μg/mL gentamicin,g/mL gentamicin, and incubated for another hour and a half. At this point, supernatant samples for invasion assays and ELISA were taken. For measurement of gene expression *in vivo*, cultures were washed twice with sterile PBS, covered with DMEM with gentamicin (16 μg/mL gentamicin,g/mL), and incubated for additional 22 h. For cell invasion assays, then infected cells were lysed with 1% Triton X-100–PBS (pH 7.4) (10 min, room temperature), and the number of viable intracellular bacteria was determined by plating of appropriate dilutions in LB agar, and colony counting.

### Competitive proliferation assays within RAW murine macrophages

RAW264.7 (murine macrophages; ECACC no. 91062702) cells were seeded in 24-well plates at 1.5 × 10^5^ cells per well and incubated in DMEM supplemented with 10% fetal calf serum for 24 h at 37 °C with 5% CO2. Bacterial strains were streaked individually onto LB agar plates supplemented with the appropriate antibiotic, and glass tubes containing LB were inoculated with each strain, incubated overnight at 37 °C with shaking, and mixtures of different genotypes at 1:1 ratio were added to eucaryotic cells cultured to up to 60 to 70% confluency in 24-well plastic plates (approximately 3 x 10^5^ eucaryotic cells/well) at a multiplicity of infection of 50. To remove noninternalized bacteria, cell cultures were washed twice with sterile PBS 30 minutes after infection, overlaid with DMEM containing 100 μg/mL gentamicin,g/mL gentamicin, and incubated for another hour and a half. The culture was then washed twice with sterile PBS, covered with DMEM with gentamicin (16 μg/mL gentamicin,g/mL), and incubated for 22 h. Then infected cells were lysed with 1% Triton X-100–PBS (pH 7.4) (10 min, room temperature), and the number of viable intracellular bacteria was determined by plating of appropriate dilutions in LB agar, and colony counting. Calculation of competitive index was done as described elsewhere^59^.

### Detection of IL-8 secretion by ELISA

Supernatants of HeLa human epithelial cells either non-infected or infected by the different bacterial genotypes, were assayed for IL-8 secretion using the Human IL-8 Uncoated ELISA kit (Invitrogen), following manufacturer’s instructions. Briefly, 96-well polysorbent plates previously incubated overnight at room temperature with human IL-8 capture antibody were washed, and then blocked with Reagent Diluent overnight at 4°C, and washed again. Then, 100 µL of appropriately diluted samples of supernatant (and serial dilutions of an appropriate human IL-8 standard, for IL-8 quantification) were transferred to the plates and incubated for 2 h at room temperature. Then, plates were washed again, followed by 2 h of incubation at room temperature with IL-8 detection antibody, a new wash, and 20 min of incubation, at room temperature and in the dark, with streptavidin-HRP B. Plates were then washed one final time, and incubated for 20 min at room temperature in the dark with substrate solution, upon which STOP solution was added, and signal development and detection were carried out using a Biotek SYNERGY HTX® multi-mode benchtop microplate reader (Agilent, Santa Clara, CA, United States of America), at a wavelength of 450 nm, with a subtraction wavelength of 570 nm. Serially diluted IL-8 standards were used for the extrapolation of protein concentration from absorbance data.

### Viability spot assays at neutral and low pH

For viability assays at normal and low pH, strains were first streaked out of their respective frozen vials, then individual colonies were inoculated separately in medium without antibiotics and incubated overnight (approximately 16 h); the next morning, 10^-1^, 10^-2^, 10^-3^, 10^-4^, 10^-5^, 10^-6^ and 10^-7^ dilutions were prepared by serial dilutions in sterile PBS. Then 5 µl aliquots of these bacterial solutions (approximately 5 × 10^6^ to 0.5 cells) were spotted onto plain LB agar plates at pHs of either 6.8 or 4.9, and these plates were incubated overnight at 37 °C.

### Statistical analyses

All analyses were conducted using Libreoffice Calc version 6.0.7.3 (www.libreoffice.org) software, R version 3.6.3 (www.r-project.org) software via RStudio version 1.2.1578 interface (www.rstudio.com), and GraphPad Prism version 8.0.2 (www.graphpad.com). In competitive fitness and luminiscence measurement assays, for each set of competitions, one-sample two-tailed Student’s t-tests were used to determine whether the data set corresponding to each genotype is significantly different than 0. For comparing different conditions or genotypes, two-tailed unpaired Student’s t-tests were used.

## ACKNOWLEDGEMENTS

We are grateful to Modesto Carballo and Laura Navarro of the “Servicio de Biología” of the “Centro de Investigación, Tecnología e Innovación de la Universidad de Sevilla (CITIUS)”, for their assistance with experiments performed at CITIUS facilities.

## AUTHOR CONTRIBUTIONS

Conceptualization: R.B., F.R.-M., J.B.-B.; validation: R.B., J.B.-B., F.R.-M.; formal analysis: R.B., J.B.-B., F.R.-M; investigation: R.B., D.A.P.-D., J.J.-E., M.O.-P; resources: F.R.-M., J.B.-B., R.B.; writing—original draft preparation: R.B.; writing—review and editing: R.B., D.A.P.-D., J.J.-E., M.O.-P, J.B.-B., F.R.-M.; supervision: F.R.-M., J.B.-B., R.B.; project administration: F.R.-M., J.B.-B., R.B.; funding acquisition, F.R.-M., J.B.-B., R.B. All authors have read and agreed to the published version of the manuscript.

## SUPPLEMENTARY FIGURES

**Figure S1.**
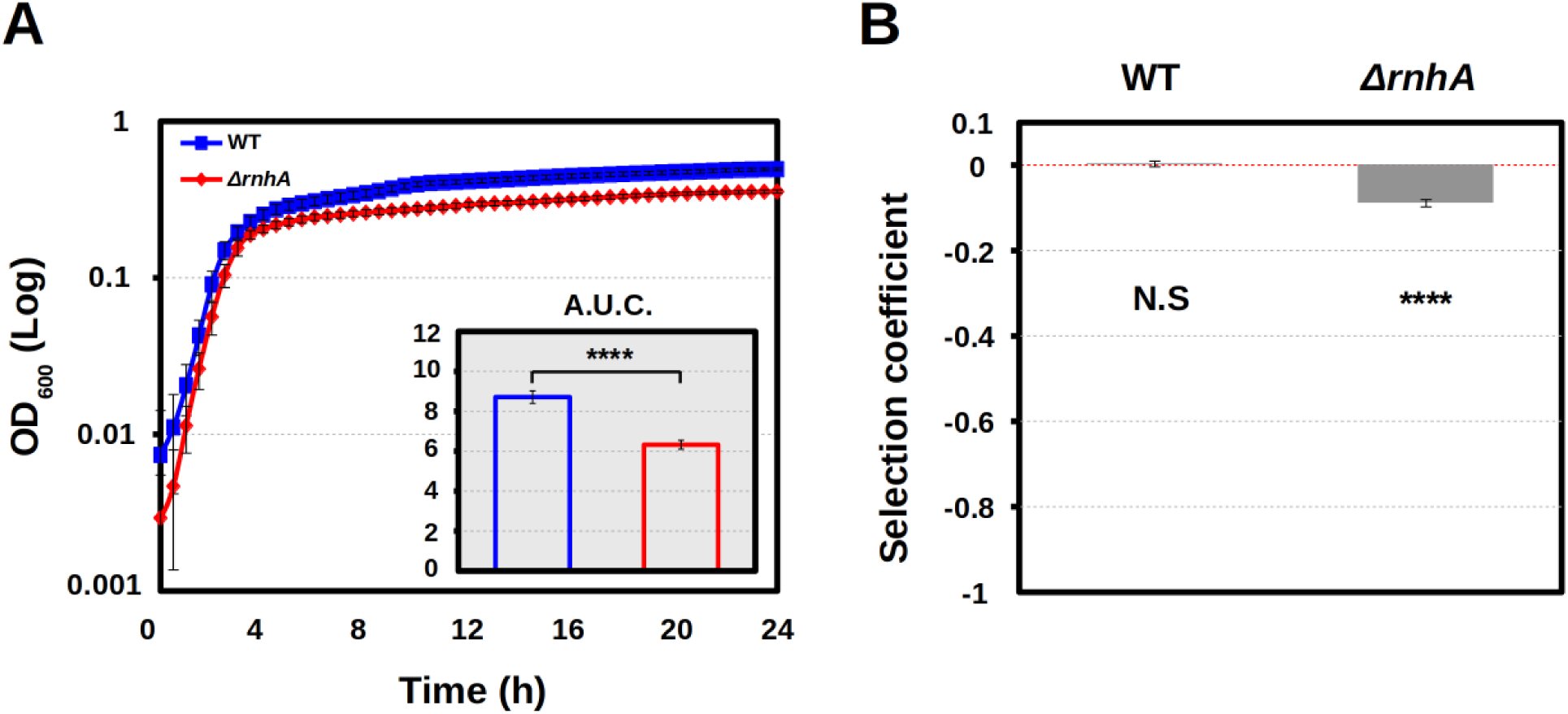
Fitness of the *ΔrnhArnhA* mutant compared to wild-type bacteria. **A.** Growth curves of wild-type (blue dots and line) and *ΔrnhA* mutant (red dots and line) bacteria in LB. Dots represent the average and error bars represent standard deviation (n ≥ 7 independent biological replicates, with two technical replicates each). **Inset:** Area Under the Curve (AUC) of wild-type (blue bar) and *ΔrnhA* mutant (red bar) in the growth curves shown. Bars represent average and error bars standard deviation (n ≥ 8 independent biological replicates, with two technical replicates each). N.S. non-significant; *P < 0.05; **P < 0.01; ***P < 0.001; ****P < 0.0001 (one-sample two-tailed Student’s t test). **B.** Competitive fitness of wild-type and *ΔrnhA* mutant bacteria against a labeled wild-type reference strain in LB. Bars represent average and error bars standard deviation (n ≥ 3 independent biological replicates). N.S. non-significant; *P < 0.05; **P < 0.01; ***P < 0.001; ****P < 0.0001 (one-sample two-tailed Student’s t test).

**Figure S2.**
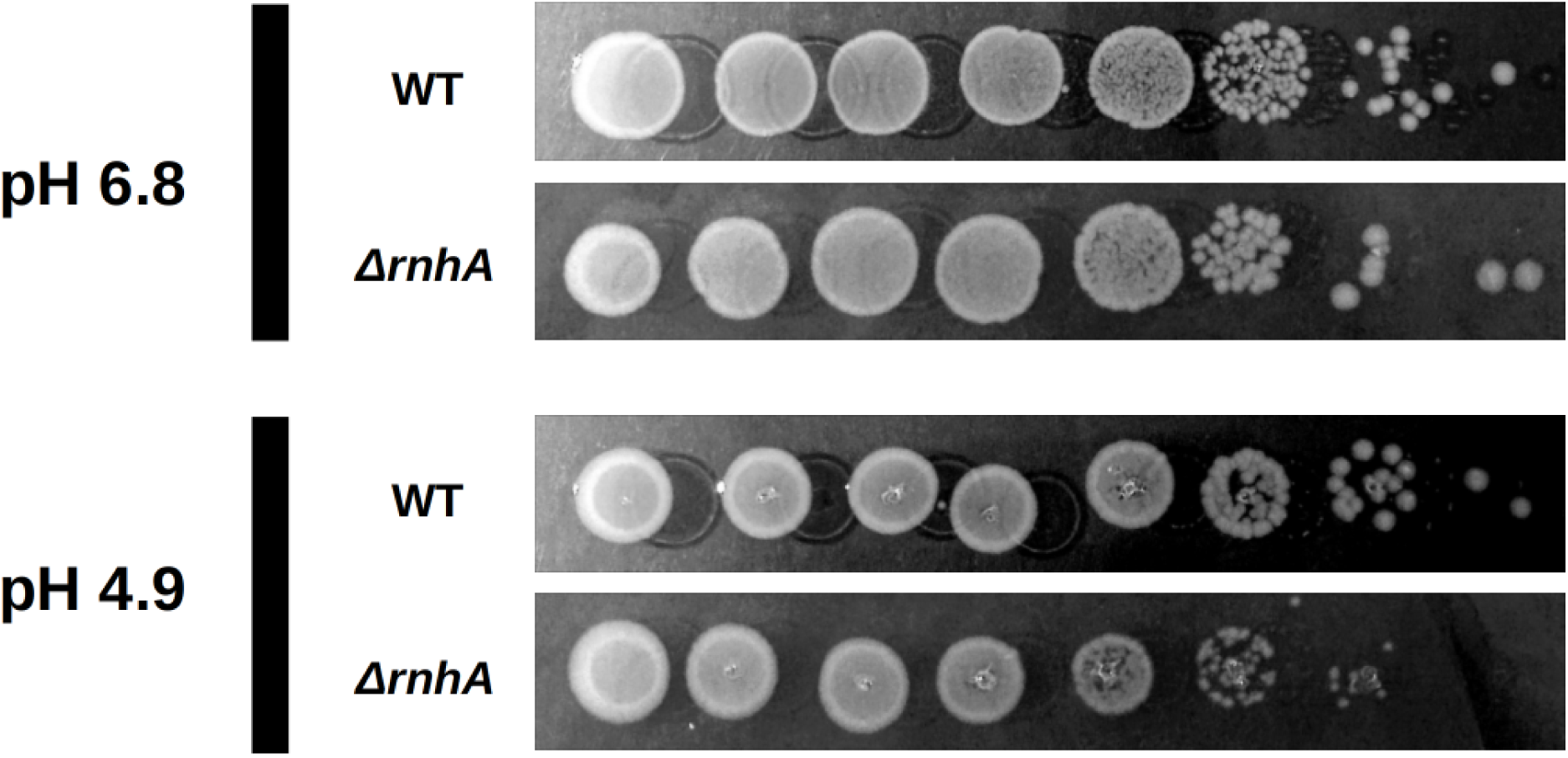
Lack of RNase HI increases sensitivity to low pH. Serially diluted wild-type (top lanes) and *ΔrnhA* (bottom lanes) in LB at pH 6.8 (top panel) or 4.9 (bottom panel). The experiment included four biological replicates and four technical replicates; representative datasets are shown.

